# The modifying enzyme Tsr3 establishes the hierarchy of Rio kinase activity in 40S ribosome assembly

**DOI:** 10.1101/2021.09.28.462141

**Authors:** Haina Huang, Melissa Parker, Katrin Karbstein

## Abstract

Ribosome assembly is an intricate process, which in eukaryotes is promoted by a large machinery comprised of over 200 assembly factors (AF) that enable the modification, folding, and processing of the ribosomal RNA (rRNA) and the binding of the 79 ribosomal proteins. While some early assembly steps occur via parallel pathways, the process overall is highly hierarchical, which allows for the integration of maturation steps with quality control processes that ensure only fully and correctly assembled subunits are released into the translating pool. How exactly this hierarchy is established, in particular given that there are many instances of RNA substrate “handover” from one highly related AF to another remains to be determined. Here we have investigated the role of Tsr3, which installs a universally conserved modification in the P-site of the small ribosomal subunit late in assembly. Our data demonstrate that Tsr3 separates the activities of the Rio kinases, Rio2 and Rio1, with whom it shares a binding site. By binding after Rio2 dissociation, Tsr3 prevents rebinding of Rio2, promoting forward assembly. After rRNA modification is complete, Tsr3 dissociates, thereby allowing for recruitment of Rio1. Inactive Tsr3 blocks Rio1, which can be rescued using mutants that bypass the requirement for Rio1 activity. Finally, yeast strains lacking Tsr3 randomize the binding of the two kinases, leading to the release of immature ribosomes into the translating pool. These data demonstrate a role for Tsr3 and its modification activity in establishing a hierarchy for the function of the Rio kinases.

## Introduction

Ribosomes are molecular machines composed of 4 ribosomal RNAs (rRNAs) and 79 ribosomal proteins. Assembly of these complexes occurs in a highly coordinated manner, that starts co-transcriptionally in the nucleolus, and is completed in the cytoplasm. This process is assisted by over 200 assembly factors (AFs), which transiently bind the nascent subunits to facilitate the modification, processing and folding of the rRNAs, promote the incorporation of ribosomal proteins, and allow for regulation and quality control (Bassler and Hurt, 2019; Chaker-Margot, 2018; de la Cruz et al., 2015; Klinge and Woolford, 2019; Peña et al., 2017; Woolford and Baserga, 2013).

Structural and biochemical studies of ribosome assembly have yielded substantial insight into these molecular events. While parallel assembly pathways appear to play a role during very early stages of assembly (Cheng et al., 2019; Cheng et al., 2020; Davis et al., 2016; Du et al., 2020; Mulder et al., 2010; Sanghai et al., 2018; Sashital et al., 2014), the process overall is highly hierarchical. *E*.*g*. during the latest, cytoplasmic stages of assembly, the remaining AFs dissociate in a highly ordered manner (Ghalei et al., 2015; Ghalei et al., 2017; Huang et al., 2020; Parker et al., 2019). This hierarchy allows for the integration of quality control with progress in the assembly cascade and is thus critical for faithful assembly. How this hierarchy is achieved is not well understood.

As part of the hierarchical assembly pipeline, there are numerous instances where the same site is bound by a succession of AFs: *E*.*g*., Krr1 and Pno1 bind the same location on the platform in early and late intermediates (Sturm et al., 2017); Bms1 and Tsr1 occupy the same site on the body in early and late intermediates (Gelperin et al., 2001; Kornprobst et al., 2016; McCaughan et al., 2016); and finally, the kinases Rio2 and Rio1 share a binding site on the head in late and very late intermediates (Ameismeier et al., 2020; Strunk et al., 2011). Further complicating the establishment of hierarchy, both Krr1 and Pno1 harbor two conserved KH domains, and a nearly identical overall structure (Sturm et al., 2017); Tsr1 is an inactive structural mimic of the GTPase Bms1 (Gelperin et al., 2001; McCaughan et al., 2016; Strunk et al., 2011) and Rio1 and Rio2 share features of the RIO kinase domain (Knuppel et al., 2018). How the order in which these factors bind (and thus function) is established remains unknown. This is especially important for factors that release others, like Rio1, which dissociates Nob1 and Pno1 (Ameismeier et al., 2020; Parker et al., 2019; Plassart et al., 2021; Turowski et al., 2014). Premature binding of Rio1 (prior to Rio2) would lead to release of immature rRNA into the translating pool, which changes the proteome. Similarly, how rebinding of the earlier factor after its dissociation is avoided also remains unclear.

To address these questions, we have studied the temporal ordering in the binding and activity of the kinases Rio2 and Rio1, which are 19% identical and 43% similar, and whose structures overlap with a root mean square deviation (RMSD) of 1.6 Å (archaeal structures) or 3.4 Å (human structures) and share a binding site on the nascent subunit **(Figure 1A, Figure S1)**. Rio2 binds first, after the dissociation of the modifying enzyme Emg1, late in the nucleolar stage of assembly (**Figure 1B**). Like snR35 and Emg1, Rio2 helps chaperone the folding of the rRNA (Huang and Karbstein, 2021). Its ATPase-dependent dissociation is required for the nascent 40S subunits to enter a translation-like quality control cycle, in which maturation is coupled to a test-drive that probes the functionality of the newly made subunit (Ferreira-Cerca et al., 2012; Huang et al., 2020). At the very end of this translation like cycle, after Nob1-dependent maturation of the 3’-end of 18S rRNA, Rio1 releases both Nob1 and its binding partner Pno1, thereby allowing for the incorporation of Rps26, the final step in maturation of the 40S subunit (Ameismeier et al., 2020; Parker et al., 2019; Plassart et al., 2021; Turowski et al., 2014). How cells coordinate the activity of the two kinases to ensure that Rio2 acts before Rio1, and that Rio1 only binds after the translation-like quality control cycle is completed and not before or immediately after Rio2 remains unknown.

**Figure 1.**
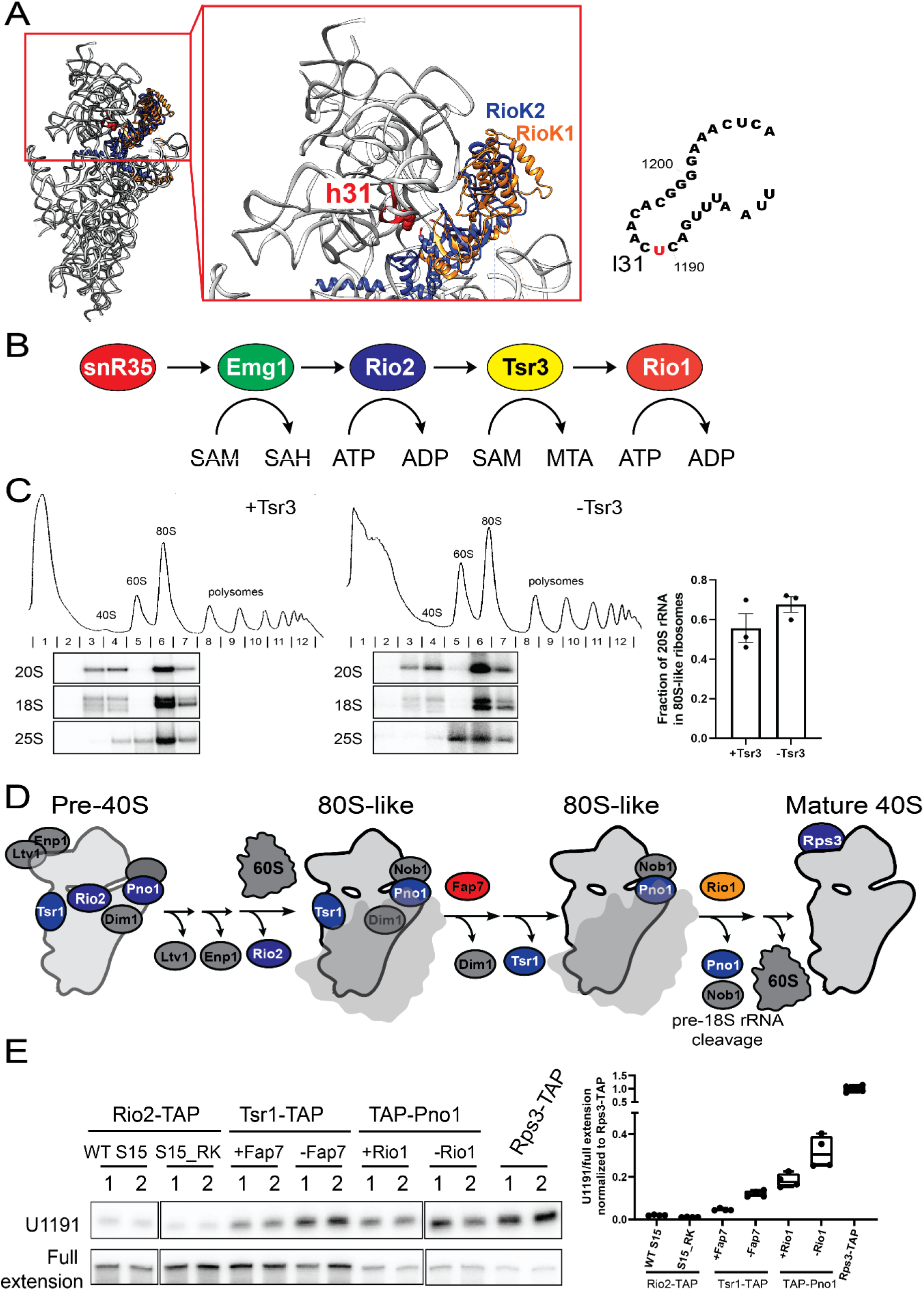
Tsr3 functions in 80S-like ribosomes. A. (Left) Superimposition of two human pre-40S structures (PDBID 6ZXE and 6G18, respectively) based on Rps15 (uS19) shows overlap of human Rio1 (RioK1, colored in orange) and human Rio2 (RioK2, colored in blue). 18S rRNA from Rio1 containing pre-40S (PDBID 6ZXE) is shown in gray ribbon, with h31 highlighted in red and U1248 (U1191 in yeast) highlighted in red sphere. (Right) Secondary structure of 18S loop 31 (l31), which is bound by all factors during assembly and contains the snR35, Emg1 and Tsr3 modification site at U1191. B. snR35, Emg1, Rio2, Tsr3 and Rio1 all bind loop 31 at successive stages of 40S maturation. Emg1 and Tsr1 utilize S-adenosyl-methionine (SAM), to produce S-adenosyl-homocysteine (SAH) or methylthioadenosine (MTA), respectively, while Rio2 and Rio1 hydrolyze ATP. C. Absorbance profiles at 254nm and corresponding northern blots of 10-50% sucrose gradients from Gal:Fap7 (+Tsr3) or ΔTsr3,Gal:Fap7 (-Tsr3) cells. In all cases Fap7 was depleted in glucose for over 16h to accumulate 80S-like ribosomes. Fractions 2-7 for each gradient were probed for 20S, 18S and 25S rRNA. Quantifications of the data on the left indicate the efficiency of formation of 80S-like ribosomes. Data are shown as mean with standard deviation (n=3). D. Simplified scheme showing ordered release of Ltv1, Enp1 and Rio2 before formation of 80S-like ribosomes, and different stages of 80S-like ribosomes facilitated by Fap7 and Rio1, respectively. Tagged proteins for pulldowns are indicated in blue. Note that Rps3 is bound in all the stages depicted here, but because mature 40S are so much more abundant, the majority of Rps3-TAP-tagged subunits are mature. E. (Left) Primer extension results indicating the installation of acp modification at U1191 in different purified pre-40S intermediates and mature 40S. Signal for U1191 was first normalized to full extension before further normalization to mature 40S (Rps3-TAP). 2 representative biological replicates (1, 2) are shown for each cell type. (Right) Quantification of the normalized U1191 stop from the left and two additional replicates (n=4).

Rio1 and Rio2 both interact with a conserved rRNA loop in the P-site, loop 31 (l31), which closes helix 31 **(Figure 1A)**. This loop is amino-carboxypropyl (acp)-modified at U1191 by the conserved modifying enzyme Tsr3 (Meyer et al., 2016). Thus, Tsr3 shares the same rRNA binding site as both Rio2 and Rio1. Here, using structural, biochemical and genetic tools, we show how Tsr3 separates the activity of the Rio2 and Rio1 kinases and helps ensure their sequential function. We first pinpoint the functional timing of Tsr3; after Rio2 dissociates, but before Rio1 associates. Genetic interactions strongly suggest a role for Tsr3 in preventing rebinding of Rio2 to ensure its release is irreversible. Our data also reveal that the modification activity of Tsr3 is required for its release from pre-40S intermediates, and that its dissociation is required for Rio1 binding. Deletion of Tsr3 leads to release of immature pre-18S rRNA into the polysomes, demonstrating the importance of Tsr3 in regulating the timing of Rio1 function.

## Results

### Tsr3 functions in 80S-like ribosomes

Immunofluorescence experiments indicate that Tsr3 resides mostly in the cytosol (Breker et al., 2014; Breker et al., 2013) (A localization atlas of the yeast proteome is available at http://www.weizmann.ac.il/molgen/loqate/). Consistently, deletion of Tsr3 leads to accumulation of the cytoplasmic pre-18S rRNA precursor, 20S rRNA(Meyer et al., 2016). During the cytoplasmic stages of 40S ribosome maturation, a translation-like cycle serves as a mechanism for testing functionality of nascent 40S subunits (Lebaron et al., 2012; Strunk et al., 2012). Specifically, pre-40S intermediates are joined by mature 60S with the help of the translation initiation factor, eIF5B. The resulting 80S-like ribosomes do not produce proteins but serve as a hub for quality control and rRNA maturation. After that, 80S-like ribosomes are broken apart by the translation termination factor Rli1. Previous work indicated that Tsr3 acted around the time of formation of 80S-like ribosomes, as modification was not observed in assembly intermediates stalled just before formation of 80S-like ribosomes. Moreover, intermediates associated with eIF5B are the first to show the modification (Hector et al., 2014).

To more accurately pinpoint the timing of Tsr3 activity during 40S maturation, we first tested whether Tsr3 was required for the formation of 80S-like ribosomes. In order to answer this question, we used a previously described *in vivo* assay, in which we stall the translation-like cycle after the formation of 80S-like ribosomes by depleting the ATPase Fap7 (Huang et al., 2020; Strunk et al., 2012). The efficiency of 80S-like ribosome formation is measured by quantifying the fraction of 20S rRNA in 80S-like ribosomes. If a protein of interest is required for the formation of 80S-like ribosomes, then the fraction of 20S rRNA found in 80S-like ribosomes decreases, as the nascent subunits are stalled in 40S-sized intermediates. The fraction of 20S rRNA in 80S-like ribosomes does not decrease in Tsr3-deletion cells relative to WT cells **(Figure 1C**). To the contrary, we observed a small increase in 80S-like ribosome formation. Together these data demonstrate that deletion of Tsr3 does not interfere with 80S-like ribosome formation and suggest that Tsr3 functions within 80S-like ribosomes, such that its deletion exacerbates the Fap7-dependent block within 80S-like ribosomes.

We further dissected Tsr3’s functional timing by asking when in the translation-like cycle its substrate is modified. As shown in the simplified scheme in **Figure 1D**, formation of 80S-like ribosomes requires the ordered release of the AFs Ltv1, Enp1 and Rio2 (Huang et al., 2020). Tsr1 and Dim1 dissociate from 80S-like ribosomes, with the ATPase Fap7 releasing Dim1 (Ghalei et al., 2017). Finally, Pno1 and Nob1 are released by the kinase Rio1 after formation of mature 18S rRNA (Ameismeier et al., 2020; Parker et al., 2019; Plassart et al., 2021; Turowski et al., 2014). Guided by this scheme, we stalled assembly at distinct stages by depletion and mutation of these AFs and then used TAP-tags on different assembly factors to isolate intermediates at different stages of the translation-like cycle. This is necessary to enrich the desired intermediates: *e*.*g*. Rio2 cannot pull down 80S-like ribosomes because it dissociates prior to their formation.

To stall assembly immediately prior to release of Rio2 and the formation of 80S-like ribosomes, we utilized a previously described mutant in Rps15/uS19, S15_RK (R137E; K142E,(Huang et al., 2020)), and isolated the intermediate using Rio2-TAP. Importantly, control experiments demonstrate that this mutation does not interfere with acp modification in mature 40S subunits **(Figure S2)**. To accumulate 80S-like intermediates prior to the release of Tsr1 and Dim1, we used Tsr1-TAP, combined with depletion of Fap7, as previously described (Ghalei et al., 2017; Rai et al., 2021). Finally, to isolate assembly intermediates immediately prior to the release of Pno1 and Nob1, we depleted Rio1, and isolated the intermediates using a TAP-tag on Pno1, as also previously described (Parker et al., 2019). To purify mature 40S subunits, we utilized S3-TAP. The presence of the Tsr3-directed modification at U1191 leads to a stall during reverse transcription, which can be visualized as a prominent band in primer extension experiments (Meyer et al., 2016).

Intermediates stalled prior to Rio2 release and the formation of 80S-like ribosomes (Rio2-TAP), do not show a substantial extension stop at U1191 (**Figure 1E**), with and without the Rps15_RK mutation. Partial modification is observed in 80S-like ribosomes stalled prior to Fap7-dependent release of Dim1, which is further increased in intermediates stalled prior to Rio1 activity **(Figure 1D)**. Together, these data strongly suggest that the modification occurs after Fap7-mediated release of Dim1 and before Rio1-dependent dissociation of Pno1 and Nob1. Thus, Tsr3 is recruited to nascent 40S ribosomes after Rio2 is released but before Rio1 is recruited. We suggest that the modification efficiency in the Rio1 depleted strain is less than in mature 40S because not all intermediates are stalled prior to the Rio1-dependent step due to an “backup” in the assembly line.

### The Rps15 C-terminal tail plays a direct role in recruiting Tsr3

To visualize how Tsr3 might be recruited to pre-ribosomes, and gain insight into its potential interacting partner(s), we manually docked the Tsr3 crystal structure onto 80S-like ribosomes, placing its substrate binding pocket and the RNA binding surface (Meyer et al., 2016) towards its rRNA substrate, U1191 **(Figure 2A)**. This dock suggests that the C-terminal extension of Rps15/uS19 might interact with Tsr3. We therefore tested whether mutations in the C-terminal tail of Rps15 affect acp modification at U1191. Using the primer extension assay described above on mature ribosomes, we found a substantial defect in acp modification in ribosomes from yeast containing the Rps15 mutant S15_YRR (S15_Y123I, R127E, R130E) **(Figure 2B**), as predicted if the C-terminal tail of Rps15 helps recruit Tsr3, similar to the docked model in **Figure 2A**. In contrast, the Rps15_RK mutation does not affect Tsr3 modification **(Figure S2)**. If the reduced acp-modification in the Rps15_YRR strain is due to defective Tsr3 recruitment, then we predict that overexpression of Tsr3 should rescue acp modification. Indeed, the modification defect introduced by S15_YRR can be almost entirely rescued by overexpression of Tsr3 **(Figure 2B)**, indicating a direct role for the Rps15 C-terminal tail in recruiting Tsr3.

**Figure 2.**
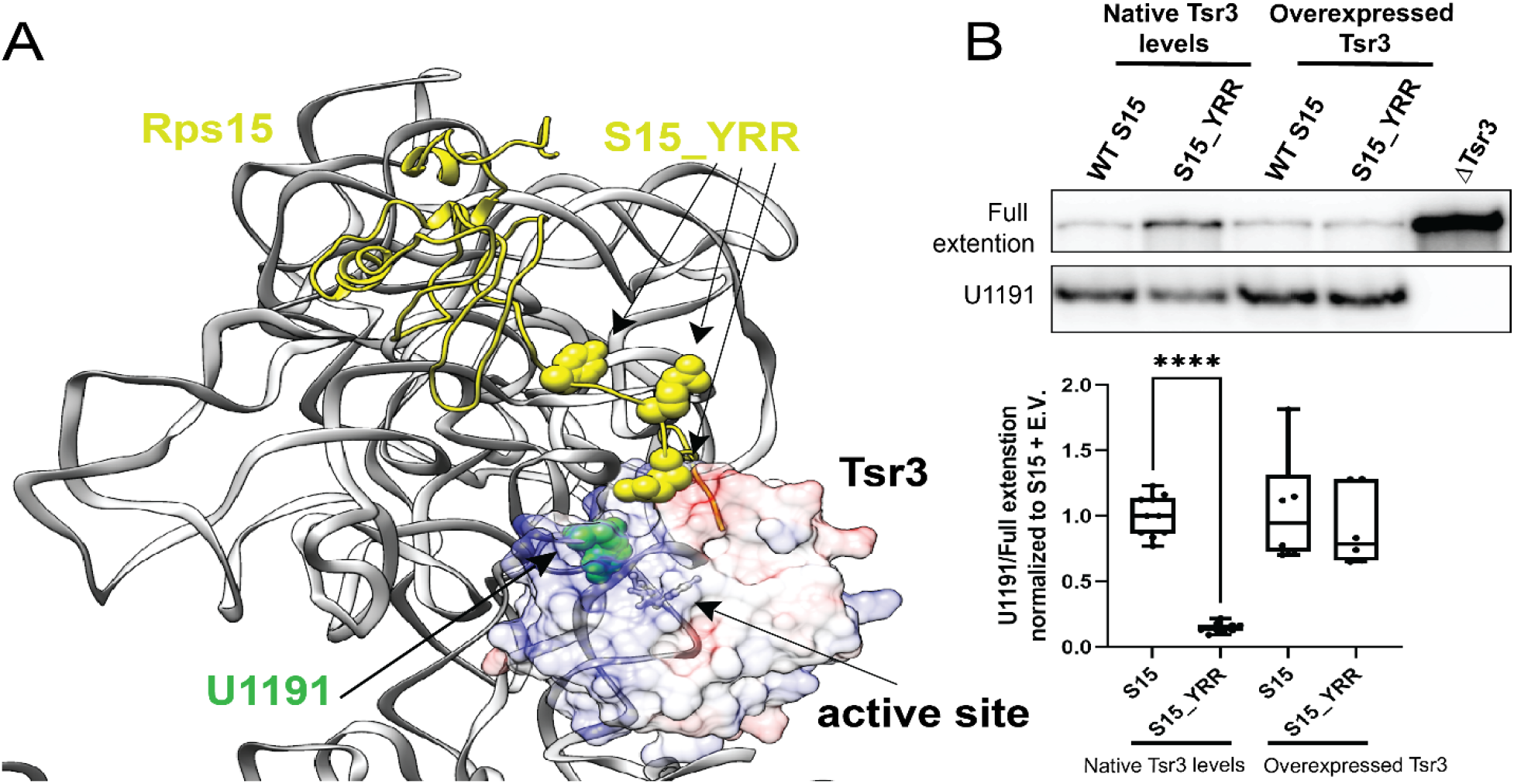
The Rps15 C-terminal tail helps recruit Tsr3. A. Predicted binding site of Tsr3 on 80S-like ribosomes. Pre-40S (PDBID: 6ZXE) and 80S-like (PDBID: 6WDR) ribosome structures were first overlayed based on 18S rRNA. The crystal structure of Tsr3 (PDBID: 5APG) was aligned to Rio1 from pre-40S (PDBID: 6ZXE) and then rotated so that its positively charged surface and its substrate binding pocket face its rRNA substrate, U1191. 18S rRNA from 80S-like ribosome (PDBID: 6WDR) is shown in gray ribbon, with U1191 highlighted in green sphere. Rps15 from 80S-like ribosome is shown in yellow, with residues, Y123, R127 and R130 highlighted in sphere. Tsr3 is shown as surface and colored based on charge (blue: positive; red: negative). The active site of Tsr3 is indicated by a black arrow based on its bound substrate analog, Se-adenosyl-selenomethionine. B. (Top) Primer extension results show acp modification at U1191 of ribosomes from Gal:S15 cells supplemented with WT S15 or S15_YRR plasmids and empty vector (E.V.) or Tsr3 plasmid. (Bottom) Quantifications of the primer extension stop at U1191 relative to the full extension from the top and additional replicates. Significance was tested using one-way ANOVA (Sidak’s multiple comparisons test). ****, p_adj_<0.0001. n≥ 6.

### Tsr3 prevents rebinding of Rio2 to ensure its release is irreversible

Above, we have shown that Tsr3 is active after Rio2 release and prior to Rio1 recruitment. Moreover, all three enzymes interact with the loop at the end of helix 31 (h31), which contains U1191 (Ameismeier et al., 2018; Ameismeier et al., 2020; Heuer et al., 2017; Meyer et al., 2016; Scaiola et al., 2018). Thus, each of these enzymes should block the binding of the other two, and Tsr3 might help order the activities of Rio2 and Rio1, by preventing the re-binding of Rio2, and blocking the premature recruitment of Rio1. To test the first part of this hypothesis, that Tsr3 blocks re-binding of Rio2, we carried out a systematic genetic screen with previously described mutants around h31 that perturb different aspects of 40S head assembly **(Figure S3A**), looking for mutants that interact genetically with Tsr3 deletion. Deletion of Tsr3 does not produce a significant growth defect in yeast **(Figure 4A**, see also (Meyer et al., 2016)) or human cells (Babaian et al., 2020). Most of the tested mutants do not show genetic interactions with Tsr3 deletion **(Figure S3B** and data not shown). In contrast, Rio2_Δloop, Rio2_K105E, S20_RK, S20_Δloop, S15_YRR and S15_RK are all synthetically sick with Tsr3 deletion **(Figure 3A)**. These mutants block the formation of 80S-like ribosomes at distinct steps **(Figure 3B**, (Huang et al., 2020)) but have two features in common. First, they all surround rRNA h31, which contains U1191, the target of Tsr3 **(Figure 3C)**. Furthermore, each of these mutants is sensitive to Rio2 overexpression **(Figure 3D & Figure S3C)**, and impacts Rio2 release (Huang et al., 2020), either directly (S15_YRR, S15_RK), or indirectly by blocking its phosphorylation, which is required for its release (Rio2_Δloop, Rio2_K105E, S20_RK, S20_Δloop). Together, these data show that Tsr3 deletion renders cells sensitive to defects in Rio2 release.

**Figure 3.**
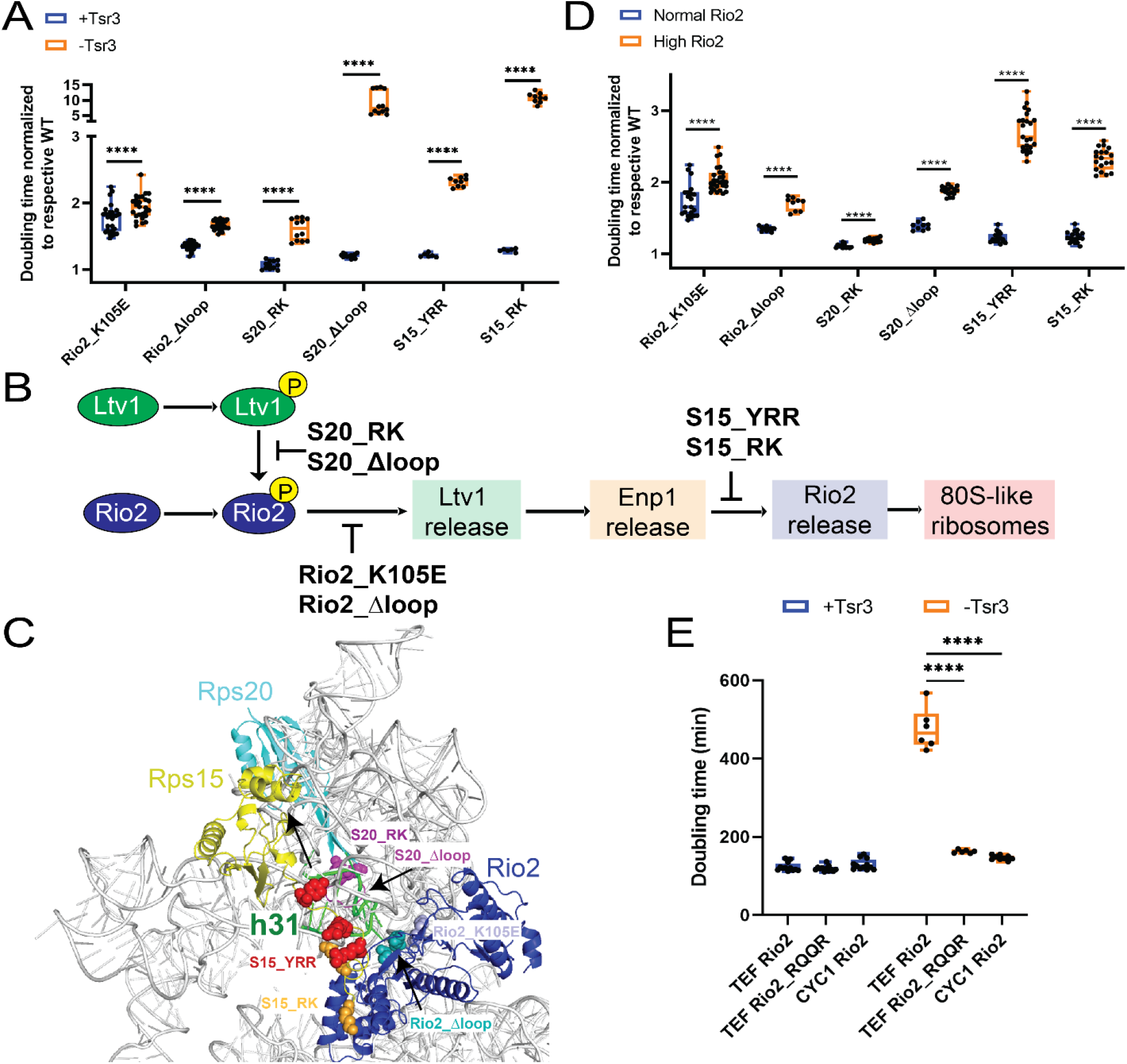
Tsr3 ensures that Rio2 release is irreversible. A. Normalized doubling times of yeast cells encoding WT Rio2/Rio2_K105E/Rio2_Δloop, WT S20/S20_RK/S20_Δloop, WT S15/S15_YRR/S15_RK in the presence and absence of Tsr3. Shown in the figure are doubling times of mutants normalized to corresponding WT protein in the same Tsr3 background. Significance was tested using a two-way ANOVA test. ****, p_adj_<0.0001. n≥6. B. Scheme showing the order steps required for the formation of 80S-like ribosomes (Huang et al., 2020) and the effects of the mutants in panel A on these steps. C. A composite structure of yeast pre-40S (PDBID 6FAI) and mature 40S (PDBID 3JAQ) obtained by overlay on Rps18. Mutations in Rio2, Rps15 and Rps20 are highlighted in sphere and colored in the same color as the corresponding text. The loop containing the residues deleted in Rio2_Δloop is unresolved in all structures and the flanking residues, R129 and S145, are highlighted in sphere in cyan. l31 from mature 40S, which is not resolved in pre-40S, is highlighted in green. D. Normalized doubling times of yeast cells encoding WT Rio2/Rio2_K105E/Rio2_Δloop, WT S20/S20_RK/S20_Δloop, WT S15/S15_YRR/S15_RK in the presence of normal levels of Rio2 (expressed under the CYC1 or native promoter) or high levels of Rio2 (expressed under the TEF promoter). Shown in the figure are doubling times of mutants normalized to corresponding WT protein in the same Rio2 background. Significance was tested using a two-way ANOVA test. ****, p_adj_<0.0001. n≥9. E. Doubling times of yeast cells encoding Rio2 or Rio2_RQQR expressed from plasmids with the lower expressing CYC1 or the higher expressing TEF promoter in the presence and absence of Tsr3. Significance was tested using a two-way ANOVA test. ****, p_adj_<0.0001. n≥6.

**Figure 4.**
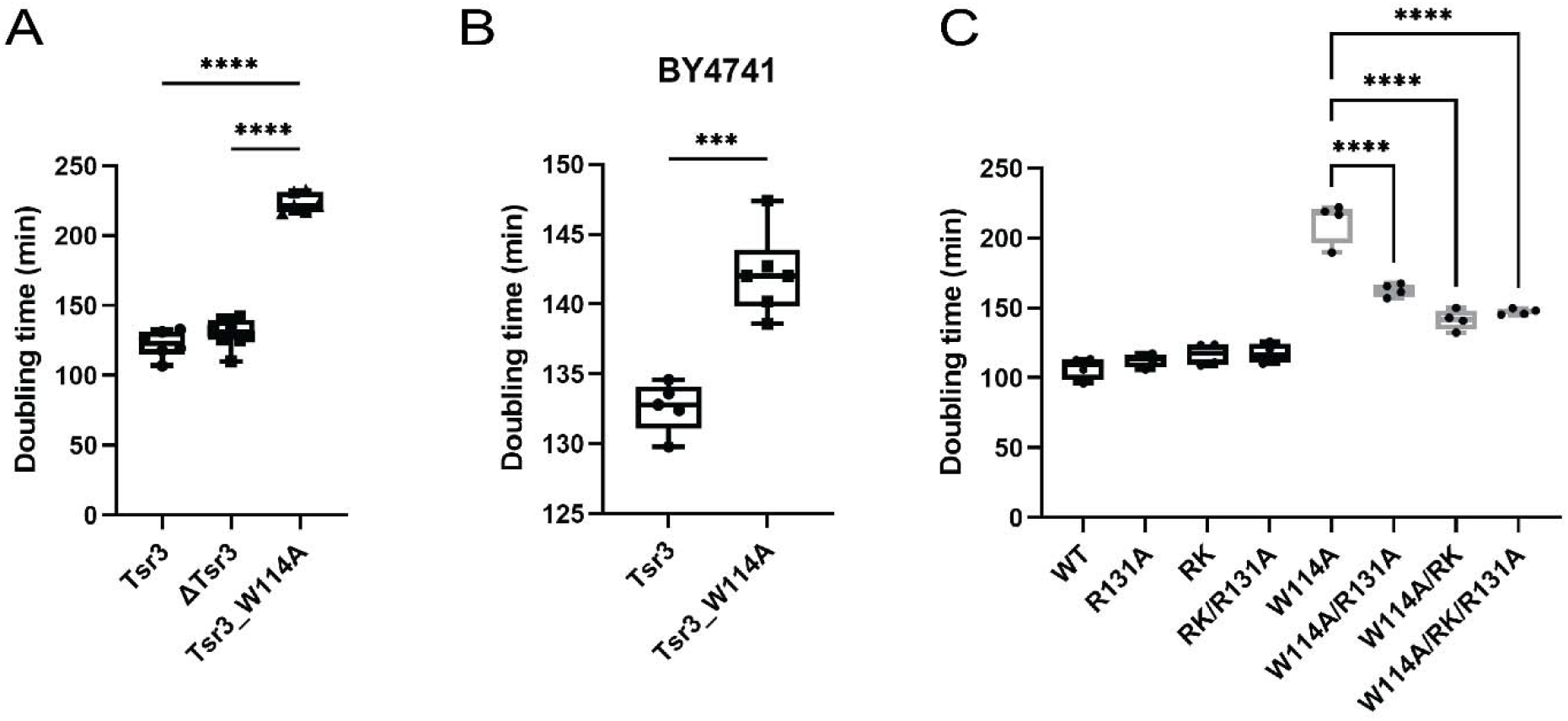
The acp modification activity of Tsr3 is required for its release. A. Doubling times of ΔTsr3 cells supplemented with a plasmid encoding WT Tsr3 (Tsr3), an empty vector (ΔTsr3) or a plasmid encoding Tsr3_W114A (Tsr3_W114A). Significance was tested using one-way ANOVA (Sidak’s multiple comparisons test). ****, p_adj_<0.0001. n≥ 6. B. Doubling times of BY4741 cells supplemented with plasmids encoding WT Tsr3 or Tsr3_W114A. Significance was tested using an unpaired t test. ***, p<0.001. n≥ 5. C. Doubling times of ΔTsr3 cells supplemented with plasmids encoding WT Tsr3 (WT), Tsr3_R131A (R131A), Tsr3_R60A,K65A (RK), Tsr3_ R60A,K65A,R131A (RK/R131A), Tsr3_W114A (W114A), Tsr3_W114A/R131A (W114A/R131A), Tsr3_W114A/R60A,K65A (W114A/RK), or Tsr3_ W114A/R60A,K65A/R131A (W114A/RK/R131A). Significance was tested using one-way ANOVA (Sidak’s multiple comparisons test). ****, p_adj_<0.0001. n≥ 4.

The observation that Tsr3 deletion renders cells sensitive to Rio2 release defects was surprising, given that Rio2 is released prior to Tsr3 binding. Nonetheless, these data can be reconciled by a model whereby Tsr3 binding prevents the reassociation of Rio2. To test this hypothesis, we tested the effect of Rio2 overexpression in the presence or absence of Tsr3. Indeed, Rio2 overexpression by the strong TEF promoter (relative to the weaker CYC1 promoter) is detrimental in ΔTsr3 cells, but not in WT cells **(Figure 3E)**. Moreover, the dominant effect from Rio2 overexpression is rescued by the Rio2 weakly-binding variant, Rio2_RQQR **(Figure 3E)**. Together, these data strongly suggest a role of Tsr3 in preventing the rebinding of Rio2, thereby ensuring its release is irreversible.

### The acp-modification activity of Tsr3 is required for its release from nascent 40S subunits

Above, we dissected the timing of Tsr3 function and its potential role in the translation-like cycle by deleting the entire protein. For many rRNA modification enzymes the presence of protein but not its modification activity is required for ribosome assembly and cell survival (Haag et al., 2015; Lafontaine et al., 1995; Meyer et al., 2011; Sharma et al., 2015; White et al., 2008; Zorbas et al., 2015). We therefore made mutations within the active site of Tsr3 to study the role of Tsr3 modification activity in ribosome assembly. We used a previously identified mutant, Tsr3_W114A, which affects the binding for the substrate, S-adenosyl-methionine (SAM), and thus impairs SAM binding and the rRNA modification activity (Meyer et al., 2016). Unexpectedly, abolishing the modification activity of Tsr3 is more detrimental to cell growth than not having Tsr3 **(Figure 4A)**. Moreover, even expressing inactive Tsr3_W114A in the background of wild type Tsr3 is detrimental **(Figure 4B)**. This dominant negative effect suggests that Tsr3’s acp-modification activity is required for its release, such that inactive protein would remain stuck on ribosomes, thereby blocking further progression of the assembly cascade. If this is true, we would predict weakly-binding variants of Tsr3 to rescue the growth defect of Tsr3_W114A. Indeed, taking advantage of a series of previously identified Tsr3 weakly-binding variants designed based on the crystal structure and validated through biochemical assays (Meyer et al., 2016), we observed a significant rescue of the growth defect from Tsr3_W114A, when this mutation was combined with any of the tested weak-binding mutations **(Figure 4C)**, further supporting the interpretation that Tsr3 activity is required for its release, and that its continued presence blocks a subsequent step(s) in 40S maturation.

### The dissociation of Tsr3 is required to accommodate Rio1 in 80S-like ribosomes

Given that Tsr3 binding is expected to be mutually exclusive not just with Rio2, but also the highly-related Rio1, which shares a binding site with Rio2 (**Figure 1A**), we hypothesized that the continued presence of Tsr3_W114A blocks the binding of Rio1, leading to the accumulation of 80S-like ribosomes and preventing the release of Nob1 and Pno1 (Parker et al., 2019).

To test if the Tsr3_W114A mutation leads to the accumulation of 80S-like ribosomes, we used gradient sedimentation to monitor whether the 18S rRNA precursor, 20S rRNA, sediments as 80S-like ribosomes or earlier 40S-sized intermediates when Tsr3 is inactivated **(Figure 5A)**. These data demonstrate that the Tsr3_W114A mutation results in the accumulation of 80S-like ribosomes (even in the presence of WT Fap7), consistent with Tsr3 blocking the activity of Rio1 (Parker et al., 2019).

**Figure 5.**
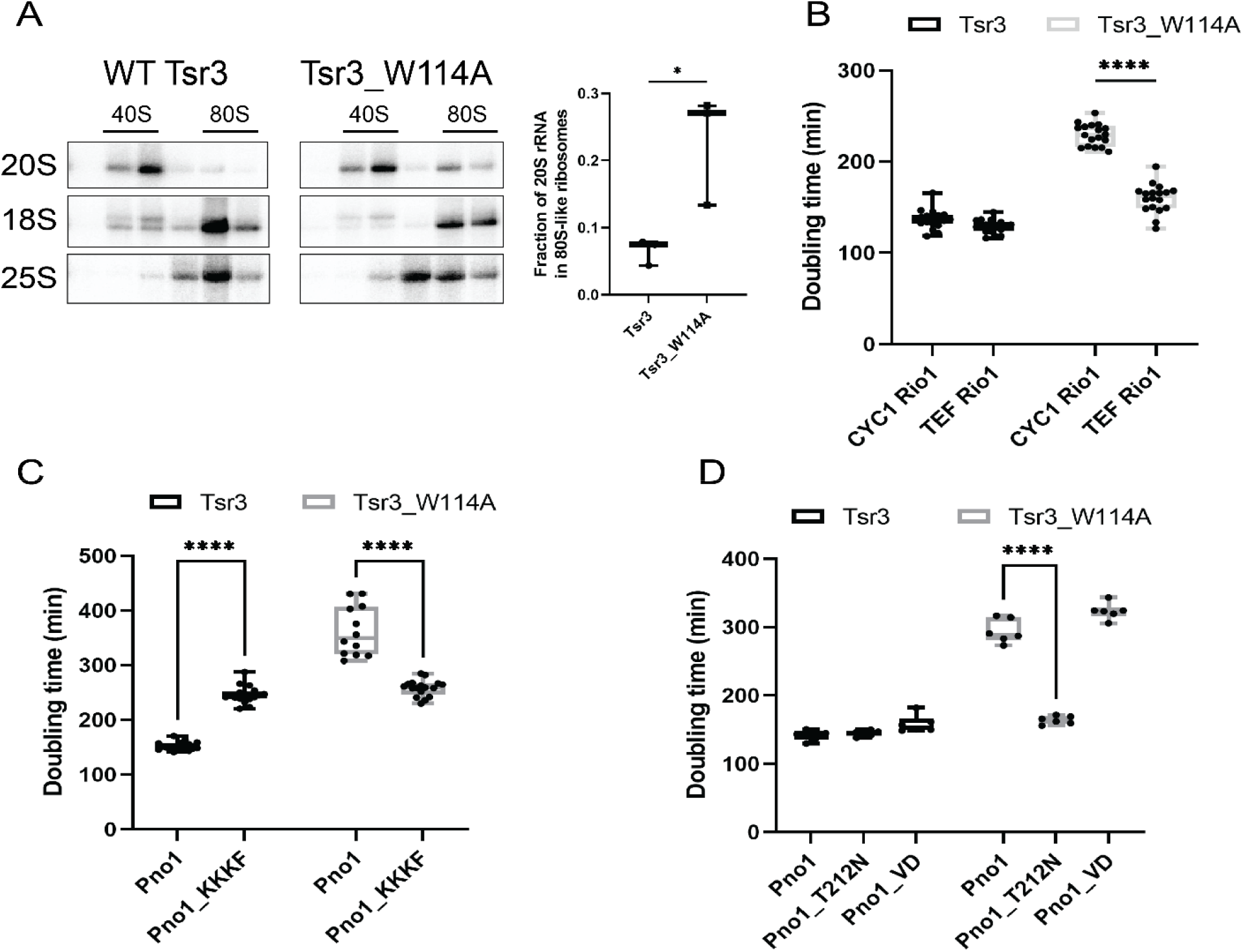
The dissociation of Tsr3 is required for Rio1 binding to pre-40S ribosomes. A. Northern blots of 10-50% sucrose gradients from ΔTsr3 cells expressing plasmid-encoded wild type (WT) HA-Tsr3 (left) or HA-Tsr3_W114A (middle) proteins. The positions where 40S and 80S ribosomes sediment are indicated. (Right) Quantifications of the data on the left indicate the efficiency of formation of 80S-like ribosome formation. Data are shown as mean with standard deviation. Significance was tested using an unpaired t test. *, p<0.05. n=3. B. Doubling times of ΔTsr3,Gal:Rio1 cells supplemented with WT Tsr3 or Tsr3_W114A and CYC1 Rio1 or TEF Rio1 plasmids. Significance was tested using a two-way ANOVA test. ****, p_adj_<0.0001. n=18. C. Doubling times of ΔTsr3,Gal:Pno1 cells supplemented with WT Tsr3 or Tsr3_W114A and Pno1 or Pno1_KKKF plasmids. Significance was tested using a two-way ANOVA test. ****, p_adj_<0.0001. n≥ 12. D. Doubling times of ΔTsr3,Gal:Pno1 cells supplemented with WT Tsr3 or Tsr3_W114A and Pno1 or Pno1_T212N or Pno1_VD plasmids. Significance was tested using a two-way ANOVA test. ****, p_adj_<0.0001. n=6.

To test the second prediction from the model that Tsr3_W114A blocks Rio1 binding, we tested if overexpressing Rio1 (under the TEF promoter) can rescue the growth defect of Tsr3_W114A. As predicted, overexpressing Rio1 almost entirely rescues the growth defect of Tsr3_W114A **(Figure 5B)**, supporting the model the Tsr3 release is required for accommodating Rio1.

To further test this model, we asked whether a previously described suppressor for the deletion of Rio1, Pno1_KKKF (K208E/K211E/K213E/F214A) (Parker et al., 2019), can rescue the growth defects observed from the Tsr3_W114A mutation. This mutation leads to weak binding of Pno1, thereby bypassing the need for Rio1 to release Pno1, and its binding partner Nob1 (Johnson et al., 2017; Parker et al., 2019). Indeed, Pno1_KKKF largely rescues the growth defect of the Tsr3_W114A mutation, even though it leads to slow growth by itself **(Figure 5C)**. This observation provides strong support for the model that Tsr3 binding blocks the recruitment of Rio1.

To further validate this suggestion, we tested whether other mutants that bypass Rio1 can rescue Tsr3_W114A. After mapping Pno1 mutations found in cancer cells (Cerami et al., 2012; Gao et al., 2013) (https://www.cbioportal.org) onto the Pno1 structure (**Figure S4A**), we focused on two mutants. Pno1_T212N is located adjacent to Pno1_KKKF, and should thus similarly weaken Pno1 binding, while Pno1_VD (V225A/D228N) appears to affect Pno1’s structure and should thus not affect its binding to ribosomes. Sucrose gradient sedimentation experiments demonstrate that as expected Pno1_T212N is bound weakly to pre-40S ribosomes, resulting in reduced co-sedimentation with ribosomes and accumulation on top of the gradient (**Figure S4C**). In contrast, Pno1_VD is not weakly bound. Moreover, Pno1_T212N, but not Rio1_VD rescues the growth defect from Rio1 depletion (**Figure S4B**), as previously shown for Pno1_KKKF (Parker et al., 2019). Thus, Pno1_KKKF and Pno1_T212N bind pre-40S weakly and rescue the growth defect from Rio1 depletion, apparently because they are self-releasing. In contrast, Pno1_VD is not self-releasing and therefore cannot rescue Rio1 depletion.

Next, we asked whether the cancer-associated Pno1_T212N also rescues the growth defect from the Tsr3_W114A mutation, as expected if this mutation blocks Rio1 binding. Indeed, Pno1_T212N can rescue the growth defects from the Tsr3_W114A mutation (**Figure 5D**). Importantly, this effect is observed only from the self-releasing Pno1_KKKF and Pno1_T212N mutations but not Pno1_VD, demonstrating that it is not a general feature of Pno1 mutants but requires the weak binding of Pno1.

### Tsr3 establishes the hierarchy of Rio kinase activity in 40S ribosome assembly

Above, we have provided evidence that the inactivation of Tsr3 impairs its release, and that the resulting persistence of Tsr3 blocks Rio1 binding. This observation suggests that the release of Tsr3 helps to temporally regulate Rio1. If this hypothesis is correct, then we predict ΔTsr3 cells to bind Rio1 prematurely, allowing for premature release of Nob1 and Pno1 and thereby the release of immature 20S rRNA into the polysomes, as previously observed in the self-releasing Pno1_KKKF (Parker et al., 2019). Indeed, we observed ≥ 2-fold increase of 20S in polysomes in Tsr3 deletion cells **(Figure 6A)**. Moreover, we also observed epistasis when ΔTsr3 is combined with Pno1_KKKF **(Figure 6B)**, which also leads to 20S escaping into polysomes. Together, these data demonstrate the importance of Tsr3-mediated temporal regulation of Rio1 binding for faithful 40S maturation.

**Figure 6.**
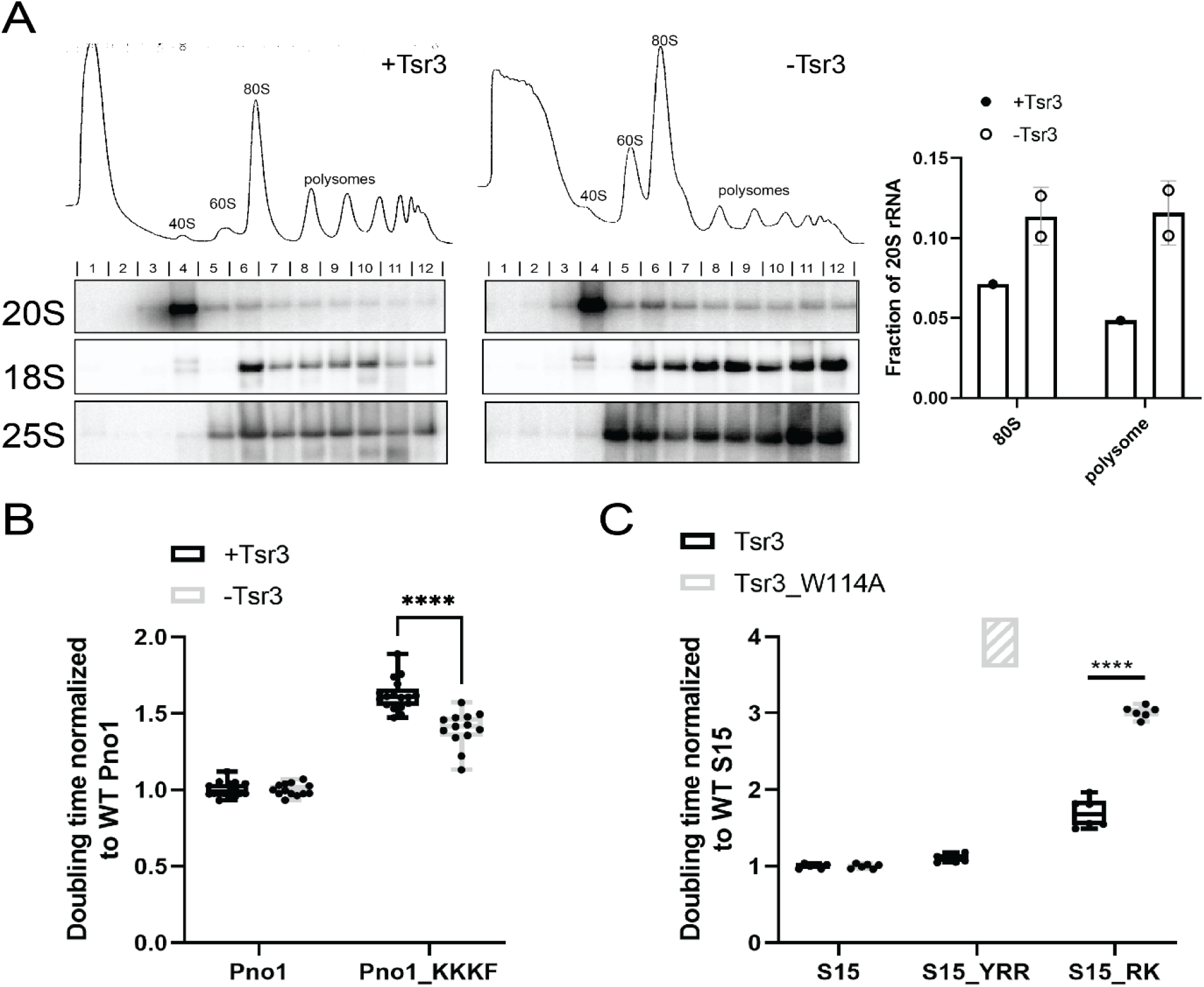
Tsr3 establishes the hierarchy of Rio kinase function. A. 10%-50% sucrose gradients of lysates from cells of WT BY4741 (+Tsr3, left) or ΔTsr3 (- Tsr3, middle). Shown below the absorbance profiles at 254 nm are Northern blots of 20S, 18S and 25S rRNA. (Right) Quantifications of the data on the left. Fraction of 20S rRNA in 80S (fraction 6-7) and polysome (fraction 8-12) are quantified. n=2. B. Normalized doubling times of ΔTsr3,Gal:Pno1 cells supplemented with WT Tsr3 or empty vector and Pno1 or Pno1_KKKF plasmids. Significance was tested using a two-way ANOVA test. ****, p_adj_<0.0001. n≥ 13. C. Normalized doubling times of ΔTsr3,Gal:S15 cells supplemented with WT Tsr3 or Tsr3_W114A and WT S15 or S15_YRR or S15_RK plasmids. Cells with Tsr3_W114A and S15_YRR plasmids do not grow and are therefore shown as a box filled with diagonal lines. Significance was tested using a two-way ANOVA test. ****, p_adj_<0.0001. n=6.

## Discussion

### Tsr3 coordinates Rio2 and Rio1 kinase activity and provides directionality to late stages of 40S maturation

Rio2 and Rio1 are two highly homologous kinases, who bind subsequent to each other to the same site on the head of the nascent small ribosomal subunit (**Figure 1A**). While there, Rio2 ensures that l31 of 18S rRNA remains unfolded, thereby chaperoning rRNA folding of the nearby junction between helices 34, 35, and 38 (Huang and Karbstein, 2021), while Rio1 releases the nuclease Nob1 and its interaction partner Pno1 after rRNA maturation is complete, thus allowing the newly-made subunit to start translation (Parker et al., 2019). How cells ensure that Rio1 functions after Rio2 and not before, which would release Nob1 from immature ribosomes and enable them to start translation, remains unclear.

Here we show that the conserved rRNA modifying enzyme Tsr3 uses its modification activity to separate and temporally organize the activities of Rio2 and Rio1, thereby ensuring that their functional order is maintained, and only correctly folded and fully matured ribosomes are released into the translating pool **(Figure 7)**.

**Figure 7.**
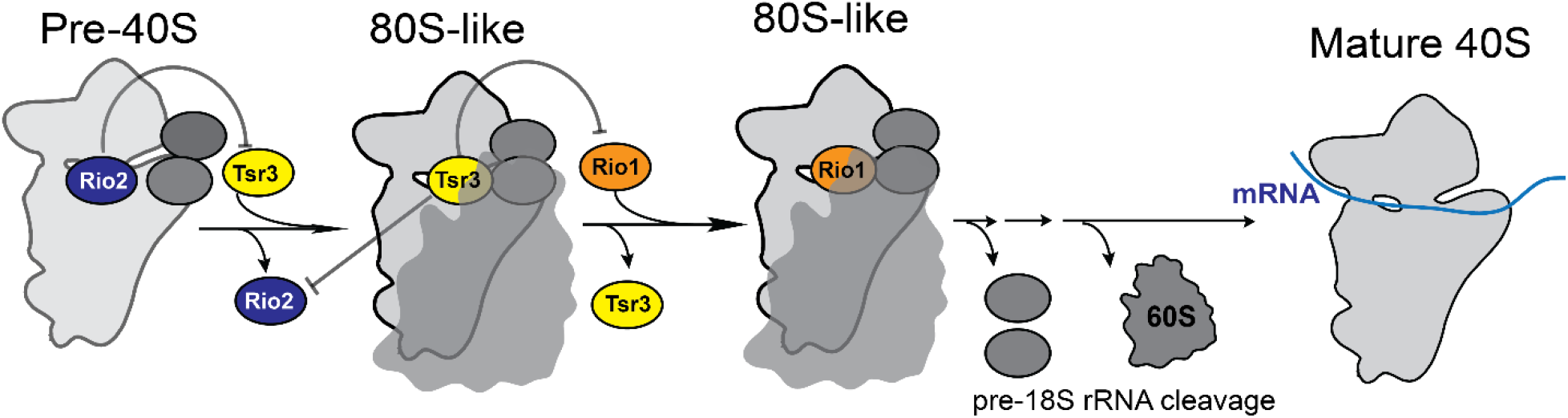
Tsr3 coordinates the activity of Rio1 and Rio2. Schematic model showing how Tsr3 coordinates and separates the activities of Rio2 and Rio1. Rio2 binding to pre-40S blocks premature binding of Tsr3. Once bound though, Tsr3 blocks re-binding of Rio2, as well as premature binding of Rio1.

Specifically, our data show that Tsr3, which shares a binding site with Rio2 and Rio1, functions within 80S-like ribosomes and thus after Rio2 is released, but before Rio1 binds. Moreover, the data also indicate that Tsr3 renders Rio2 dissociation irreversible. In addition, we show that modification activity is required to release Tsr3, and that blocking Tsr3 release blocks the binding of Rio1. Thus, together these data show that Tsr3 separates the activities of Rio2 and Rio1. The importance of this role for Tsr3 in establishing the hierarchy of binding of these two proteins is demonstrated by the observation that cells lacking Tsr3 release immature subunits into the translating pool, as expected from premature activity of Rio1.

### The ability to modify rRNA is required for Tsr3’s function in ordering Rio activity

How does Tsr3 order Rio2 and Rio1 activity, and thereby impart directionality to the process? Tsr3 utilizes S-adenosylmethionine (SAM) as a donor for the amino-carboxy-propyl- (acp) group that is transferred onto U1191 (Meyer et al., 2016), leaving behind methylthioadenosine (MTA). Because SAM is produced using ATP, Tsr3 essentially consumes energy in the form of ATP. Thus, it is possible that Tsr3 might simply utilize the energy from SAM/ATP to provide directionality. In this model, SAM-bound Tsr3 might effectively compete with both Rio2 and Rio1, while the MTA-bound Tsr3 can no longer compete with Rio1, perhaps due to some conformational differences. Additionally, or alternatively, it is also possible that the modification itself contributes to the discrimination. Indeed, the genetic interaction data support a role for Tsr3 that goes beyond simply being a competitor. Because the Tsr3_W114A mutant cannot modify rRNA but can bind the ribosomes (thereby conferring a dominant negative growth effect), we would expect it to efficiently compete against Rio2 if the role of Tsr3 was simply in competition. In this model, the Tsr3_W114A mutant should not show genetic interactions with the Rio2-release-deficient Rps15_RK or Rps15_YRR mutants. In contrast, if the ability to modify the RNA is important, then we expect the Tsr3_W114A mutant to be synthetically sick with the Rio2-release defective Rps15_YRR and Rps15_RK, as we observe for the Tsr3 deletion strain. Indeed, this is exactly what we observe (**Figure 6C)**, providing genetic evidence that the modified U1191 contributes to the discrimination against Rio2 later in assembly.

### Energy-driven changes in AF binding sites provide directionality to the process

As described above, there are numerous instances during ribosome assembly where two or more assembly factors bind the same location successively, as we have shown here for Rio2 and Rio1. Examples during small subunit assembly include snR35 and Emg1, which bind the same rRNA loop as Rio2, Tsr3 and Rio1 during the early, nucleolar stages of 40S assembly (**Figure 1B**). Similarly, the assembly factor Pno1 replaces Krr1 on the nascent platform (Sturm et al., 2017), only to be replaced with Rps26 in the final maturation step (Ferretti et al., 2017; Parker et al., 2019; Strunk et al., 2012). Moreover, Tsr1 replaces Bms1 on the nascent body (Gelperin et al., 2001; Kornprobst et al., 2016; McCaughan et al., 2016; Strunk et al., 2011). Examples during 60S maturation include Nog1, Rei1 and Reh1 binding the peptide exit channel (Fuentes et al., 2007; Greber et al., 2016; Hung and Johnson, 2006; Kallstrom et al., 2003; Klingauf-Nerurkar et al., 2020; Lebreton et al., 2006; Ma et al., 2017; Parnell and Bass, 2009; Wu et al., 2016), and Mrt4, Yvh1 and Rpp0 binding the nascent P-stalk (Kemmler et al., 2009; Lo et al., 2009). Some of these factors may also use energy for their ordered activity (Bms1 and Nog1 are GTPases and Yvh1 is annotated as a phosphatase, see also **Figure 1B**), but that is not universally true. Nonetheless, in all cases the environment of the assembly factors changes in going from the early to the late-binding factors. *E*.*g*., Tsr1 interacts with the decoding site helix, h44, while Bms1 blocks h44 incorporation into the subunit (or is blocked by its presence). In addition to being blocked by Bms1, h44 is held away by interactions with the early-binding assembly factors Utp12/Utp13 and Utp22, which appear to dissociate after maturation of the 5’-end of 18S rRNA. Thus, once te 5’-end of 18S rRNA is matured and h44 is incorporated in its native position, Bms1 will no longer be able to bind.

Similarly, while both Krr1 and Pno1 are bound very similarly to h23 and Rps14, Krr1 is also bound to Faf1, which in turn binds to the 5’-ETS helix H6. Thus, early in transcription of rRNA, Faf1 is recruited via binding to the 5’-ETS, and then can recruit Krr1, over Pno1. During assembly, H6 is degraded, while formation of h24 and h45, which is stabilized by the recruitment of Dim1, excludes Krr1 via steric contacts. In contrast, Pno1 is stabilized by an interaction with h45. Thus, the temporal ordering of Krr1 and Pno1 (or Bms1/Tsr1) binding is achieved by changes in the rRNA binding site, akin to the Tsr3-dependent changes in the Rio binding site. Notably the differences are that in the case of Krr1/Pno1 and Bms1/Tsr1, the changes arises from rRNA folding and maturation, while in the Rio case, the changes arise from a covalent modification. This might be necessary because the subunits are so far matured that few (and small) changes in folding or processing are left to impose order.

### Other modification enzymes also require their activity for release

Making the release of Tsr3 and subsequent 40S maturation steps dependent on the completion of its modification reaction ensures that most, if not all, subunits are modified. Interestingly, Tsr3 is not the only rRNA modification enzyme whose release appears to require enzymatic activity. Rrp8 and Bmt2, which methylate the N1 of A645 and A2142 in 25S rRNA, respectively, both appear to have the same property (Peifer et al., 2012; Sharma et al., 2013). Both of these enzymes are non-essential and, like Tsr3, do not demonstrate any phenotype upon their deletion (at 30 ºC). In contrast, non-functional mutants of Rrp8 and Bmt2 both lead to the accumulation of half-mer ribosomes, indicative of defects in 60S maturation (Peifer et al., 2012; Sharma et al., 2018). Like Tsr3, Rrp8 functions after snoRNA mediated rRNA modification at A649/650. Intriguingly, A649 is involved in a highly unusual RNA structure, which sets up the backend of the large subunit’s P and A-sites, again similar to Tsr3, whose target site forms the small subunit’s P-site. Furthermore, for the bacterial homolog of Dim1, KsgA, it has also been reported that its activity is required for its release, leading to strong dominant negative effects from an inactive mutant (Connolly et al., 2008). Dim1 modifies residues in the small subunit’s P-site. These defects are not observed in the yeast system (Pulicherla et al., 2009), presumably because in yeast Dim1 is actively released via the ATPase Fap7 (Ghalei et al., 2017). Thus, it appears likely that Tsr3 and Rrp8 (and bacterial KsgA) might have evolved to ensure a high modification rate in the functional centers of the ribosome.

What function does the modification of Tsr3 serve? As described above, the Tsr3 modification site forms part of the small subunit’s P-site. Yet, deletion of Tsr3 in yeast provides a less than 10% defect in growth, and no obvious defect in global translation. Similarly, Tsr3 deletion in human cells does not produce growth or global translation defects (Babaian et al., 2020). Nonetheless, a very small subset of mRNAs (<20) displays statistically significant changes in translational efficiency, although not enough to change the proteome. However, what is unique about these mRNAs, and thus causes the sensitivity to the absence of the modification, remains to be seen. Of note, many cancer cells are characterized by reduced levels of the modification (Babaian et al., 2020), supporting its importance for ribosome assembly and/or function.

## Methods

### Plasmids and yeast strains

Yeast strains **(Table S1)** were obtained from Yeast Knockout Collection (Horizon) or generated by standard homologous recombination (Longtine et al., 1998), and confirmed by PCR, serial dilution and western blotting if antibodies were available. Plasmids **(Table S2)** were constructed using standard cloning techniques and confirmed via Sanger sequencing.

### Growth curve measurements

Freshly transformed cells were grown in YPD or glucose minimal media (if the supplemented plasmid is not essential) overnight, and then diluted into fresh YPD for 3-6 hours before inoculating into 96-well plates (Thermo Scientific) at a starting OD600 between 0.04 to 0.1. A Synergy.2 plate reader (BioTek) was used to record the OD600 for 48 hours, while shaking at 30 °C. Doubling times were calculated using data points within the mid-log phase using GraphPad Prism 9. Data were averaged from at least 6 biological replicates of 3 different colonies and 2 independent measurements. For all the growth curve assays, detailed yeast strains and plasmids used in each study were summarized in **Table S3**. Statistical analyses for each measurement are detailed in the respective figure legend.

### Serial dilution

Overnight cultured cells were spotted on glucose or galactose dropout plates with 10-fold serial dilutions. Size of single colonies is used to assess cell growth.

### Primer extension assay for acp-modification

Different pre-40S intermediates were purified using TAP tagged proteins via IgG beads as previously described (Huang and Karbstein, 2021; Parker et al., 2019). Mature 40S were purified via Rps3-TAP. rRNAs from pre-40S intermediates were phenol chloroform extracted and further cleaned using RNeasy Mini columns (Qiagen). 1ug of rRNA was used for each reverse transcription reaction using a primer that starts at 1241 of 18S rRNA (sequence listed in **Table S4**). rRNA was annealed with radioactively labeled primer before reverse transcription with SuperScript III (Thermo Fisher Scientific) at 50°C for 10min. rRNAs were then hydrolyzed by addition of 200mM NaOH at 95°C for 5min. cDNAs were separated by 8% TBE-urea gels and visualized by Typhoon™ FLA 9000 (GE Healthcare). Sequencing ladders were made using the Sequenase 2.0 DNA Sequencing Kit (Thermo Fisher Scientific) and the same primer as above and 35S rDNA plasmid as template. As previously described (Meyer et al., 2016), acp modification causes a prominent stop during reverse transcription, which we validated by the sequencing ladders and the missing stop signal in ribosomes from ΔTsr3 yeast. When assaying mature ribosomes, primer extensions were carried using total RNA from respective cells. Total RNAs was isolated using the hot phenol method and cleaned by RNeasy Mini column.

### In vivo subunit joining assay

Gal:Fap7 strains in combination with the additional mutation of interest were grown to mid-log phase in galactose before inoculating into YPD for at least 16 hours to deplete Fap7 and accumulate 80S-like ribosomes. Formation of 80S-like ribosomes was assayed by sucrose gradient fractionation and Northern blotting as previously described (Huang et al., 2020; Strunk et al., 2012), and the fraction of 20S rRNA in 80S-like ribosomes quantified. Oligos used for Northern analysis are listed in **Table S4**. At least two biological replicates were assayed for each mutant.

### Quantification and statistical analysis

Northern blots were visualized by Typhoon™ FLA 9000 (GE Healthcare). Quantity One (Bio-Rad) was used to analyze Northern blot results. Western blot images were taken by imager ChemiDoc MP Imaging System from Bio-Rad after applying luminescence substrates (Bio-Rad). Image Lab (Bio-Rad) was used to analyze Western blot results. Statistical analyses were performed using GraphPad Prism version 9 (GraphPad Software, San Diego, California). Statistical tests were used as indicated in the respective figure legends.

## Acknowledgements

We thank members of the Karbstein laboratory for discussion and comments on the manuscript. This work was supported by NIH grant R35-GM136323, and HHMI Faculty Scholar grant 55108536 to K.K., and a Farris Foundation fellowship to H.H..

